# The rubber tree kinome: genome-wide characterization and insights into coexpression patterns associated with abiotic stress responses

**DOI:** 10.1101/2022.08.24.505065

**Authors:** Lucas Borges dos Santos, Alexandre Hild Aono, Felipe Roberto Francisco, Carla Cristina da Silva, Livia Moura Souza, Anete Pereira de Souza

**Author notes:** The authors contributed equally (Lucas Borges dos Santos, Alexandre Hild Aono, Felipe Roberto Francisco). Corresponding author Email addresses (Lucas Borges dos Santos), (Alexandre Hild Aono), (Felipe Roberto Francisco), (Carla Cristina da Silva), (Livia Moura Souza), (Anete Pereira de Souza).

## Abstract

The protein kinase (PK) superfamily constitutes one of the largest and most conserved protein families in eukaryotic genomes, comprising core components of signaling pathways in cell regulation. Despite its remarkable relevance, only a few kinase families have been studied in *Hevea brasiliensis*. A comprehensive characterization and global expression analysis of the PK superfamily, however, is currently lacking. In this study, with the aim of providing novel inferences about the mechanisms associated with the stress response developed by PKs and retained throughout evolution, we identified and characterized the entire set of PKs, also known as the kinome, present in the Hevea genome. A total of 1,809 PK genes were identified using the current reference genome assembly at the scaffold level, and 1,379 PK genes were identified using the latest chromosome-level assembly and combined into a single set of 2,842 PKs. These proteins were further classified into 20 different groups and 122 families, exhibiting high compositional similarities among family members and with two phylogenetically close species (*Manihot esculenta* and *Ricinus communis*). Different RNA-sequencing datasets were employed to identify tissue-specific expression patterns and potential correspondences between different rubber tree genotypes. In addition, coexpression networks under several abiotic stress conditions, such as cold, drought and latex overexploitation, were employed to elucidate associations between families and tissues/stresses. Through the joint investigation of tandemly duplicated kinases, transposable elements, gene expression patterns, and coexpression events, we provided insights into the understanding of the cell regulation mechanisms in response to several conditions, which can often lead to a significant reduction in rubber yield.

## 1. Introduction

Rubber is one of the world’s major commodities and is extensively used in various industrial and domestic applications, yielding more than 40 billion dollars annually (Board, 2018). The major source of latex for rubber production is *Hevea brasiliensis* (Hbr), commonly referred to as rubber tree, a perennial native plant from the Amazon rainforest belonging to the *Euphorbiaceae* family (Priyadarshan & Goncalves, 2003). Although the warm and humid weather in the Amazon region offers a favourable climate for Hbr growth and propagation, large-scale cultivation of Hbr is unviable due to the incidence of a highly pathogenic fungus, *Pseudocercospora ulei* (Hora Júnior et al., 2014). Thus, rubber tree plantations were transferred to other countries and regions, which could not offer optimal conditions for developing tropical crops due to the low temperatures during winter, dry periods, and elevated wind incidence (Hoa et al., 1998). Exposure to these abiotic stresses often leads to a significant reduction in latex production in most Hbr wild varieties, which has stimulated the development of breeding programs with a focus on stress-tolerant cultivars (Priyadarshan & Goncalves, 2003; Pushparajah, 1983).

Different types of abiotic stresses may trigger several physiological responses in susceptible rubber tree genotypes and often impact their survival, growth and productivity, depending on the age and vigour of the affected plant (Kuruvilla et al., 2017). In general, cold and drought stresses result in the inhibition of photosynthesis and chlorophyll degradation (Devakumar et al., 2002). Water deficit may affect the plant growth and canopy architecture of trees, and its impact during tapping seasons tends to be more severe due to the deviation of resources (carbon and water) caused by wounding stress (Devakumar et al., 1999; Kunjet et al., 2013). Cold damage leads to a decrease in membrane permeability (Meti et al., 2003; Sevillano et al., 2009), together with photosynthesis inhibition, causing more critical injuries to the plant, such as the wilting and yellowing of leaves, interveinal chlorosis, darkening of the green bark, reduction of latex flow and dieback of shoots (Meti et al., 2003).

Projections indicate that climate changes caused by global warming will enable the expansion of areas suitable for rubber tree plantation, especially in regions with greater production of natural rubber (Yang et al., 2019; Zomer et al., 2014). However, the real impact of climate change on the origin of rubber tree diversity is still unknown, but some studies suggest that these changes will have severe consequences for rubber tree biodiversity, mainly due to changes in the water regime (Marengo et al., 2018; do Prado Tanure et al., 2020). These alterations are even worse considering that the rubber tree is still being domesticated and has little genetic variability explored (De Souza et al., 2018).

The ability to sense and adapt to adverse conditions relies on the activation of complex signaling networks that protect plants from potential damage caused by these environmental changes (Kovtun et al., 2000). Protein kinases (PKs) comprise one of the most diverse protein superfamilies in eukaryotic organisms (Liu et al., 2015) and act as key components in stimulus perception and signal transduction through a chain of phosphorylation events, resulting in the activation of genes and several cellular responses (Colcombet & Hirt, 2008). The expansion of this family underlies several mechanisms of gene duplication throughout the evolutionary history of eukaryotes, including chromosomal and whole-genome duplication, tandem duplication in linked regions and retroposition events, leading to more specialized or novel gene functions (Zhang, 2003).

In the rubber tree, several kinase families have been characterized, including the mitogen-activated protein kinase (MAPK) (Jin et al., 2017), calcium-dependent protein kinase (CDPK) (Xiao et al., 2017; Zhu et al., 2018b), CDPK-related protein kinase (CPK) (Xiao et al., 2017), and sucrose non-fermenting 1-related protein kinase 2 (SnRK2) (Guo et al., 2017). These studies revealed contrasting expression patterns of the kinase families among tissues, in addition to the elevated expression of the SnRK2 and CPK families in laticifers in response to ethylene, ABA, and jasmonate stimulation (Guo et al., 2017; Zhu et al., 2018b), suggesting their potential participation during several developmental and stress-responsive processes. However, the comprehensive identification and characterization of rubber tree PKs have not yet been performed and would greatly benefit plant science research to promote a better understanding of the molecular mechanisms underlying the stress response. Furthermore, the complete characterization of the rubber tree kinome has the potential to highlight important PKs associated with stress resilience across evolution, which is of great interest for plant breeding efforts, especially considering the current genetic engineering and genome editing methodologies available (Pandita & Wani, 2021). Due to the long breeding cycles for the development of rubber tree cultivars (ranging from 25 to 30 years (Priyadarshan, 2017)) and the need for introgression of resistant alleles to more productive varieties, the definition of key molecular mechanisms signaling stress response represents an important contribution to target gene candidates for molecular breeding approaches.

In this study, we investigated the kinase diversity present in the Hbr genome through a characterization of its PKs, including the subfamily classification and the prediction of several protein properties, such as molecular weight, subcellular localization, and biological functions. The rubber tree kinome, defined as the complete repertoire of PKs, was estimated using a combined analysis with the two major genome assemblies of the rubber plant and comparative analyses with cassava (*Manihot esculenta* (Mes)) and castor plant (*Ricinus communis* (Rco)) kinomes. Furthermore, RNA sequencing (RNA-Seq) data from different Hbr genotypes were used to identify expression patterns of the kinase subfamilies, followed by the construction of gene coexpression networks for control and abiotic stress conditions. In addition to fully characterizing the Hbr kinome, our study aims to provide novel inferences about the mechanisms associated with the stress response developed by PKs and retained throughout evolution, assessed by a joint analysis considering tandemly duplicated PK subfamilies, transposable elements and expression patterns. Our study provides broad resources for future functional investigations and valuable insights into the major components associated with cell adaptation in response to environmental stresses.

## 2. Material and methods

### 2.1. Data acquisition

Sequence and annotation files of Hbr, Mes, and Rco were downloaded from the NCBI (Geer et al., 2010) and Phytozome v.13 (Goodstein et al., 2012) databases. We selected the latest genomes of cassava v.7.1 (Bredeson et al., 2016) and castor plant v.0.1 (Chan et al., 2010), as well as two major genomes of the rubber tree: the latest chromosome-level genome (Liu et al., 2020b) (Hb chr) and the reference scaffold-level assembly (Tang et al., 2016) (Hb scaf), under accession numbers PRJNA587314 and PRJNA310386 in GenBank, respectively. The same data analysis procedures for PK identification and characterization were applied to Hbr, Mes and Rco.

### 2.2. Kinome Definition

The hidden Markov models (HMMs) of the two typical kinase domains, Pkinase (PF00069) and Pkinase Tyr (PF07714), were retrieved from the Pfam database (El-Gebali et al., 2019). To select putative proteins having one or more kinase domains, protein sequences were aligned to each HMM profile using HMMER v.3.3 (Finn et al., 2011) (E-value of 1.0E-10). We retained only sequences covering at least 50% of the respective domain and the longest isoform.

The Hbr kinome was created as a combination of putative PKs identified from two different genomic datasets: Hb chr and Hb scaf. To avoid redundancy, we combined the sets using CD-HIT v.4.8.1 software (Fu et al., 2012) with the following selection criteria: (i) for proteins present in both sets as a single copy, the longest sequence was retained, and the other one was discarded; and (ii) when putative duplications were present, i.e., there were protein clusters with significant similarities in both Hb chr and Hb scaf, and all proteins from the largest set were retained. For pairwise comparisons, we set a minimum sequence identity threshold of 95% and a maximum length difference of 75%.

### 2.3. Kinase characterization and phylogenetic analyses

The PKs were classified into groups and subfamilies according to the HMMs of each family built from four plant model species (*Arabidopsis thaliana*, *Chlamydomonas reinhardtii*, *Oryza sativa*, and *Physcomitrella patens*) and supported among 21 other plant species (Lehti-Shiu & Shiu, 2012). The classification was further validated through phylogenetic analyses. The domain sequences from all PKs were aligned using Muscle v.8.31 (Edgar, 2004), and a phylogenetic tree was constructed for each kinase dataset using the maximum likelihood approach in FastTree v.2.1.10 software (Price et al., 2010) with 1,000 bootstraps and default parameters through the CIPRES gateway (Miller et al., 2011). The resulting dendrograms were visualized and plotted using the statistical software R (Ihaka & Gentleman, 1996) together with the ggtree (Yu et al., 2017) and ggplot2 (Wickham et al., 2016) packages.

For each PK, we obtained the following characteristics: (a) gene location and intron number, according to the GFF annotation files; (b) molecular weight and isoelectric point with ExPASy (Gasteiger et al., 2003); (c) subcellular localization prediction using CELLO v.2.5 (Yu et al., 2006) and LOCALIZER v.1.0.4 (Sperschneider et al., 2017) software; (d) the presence of transmembrane domains using TMHMM Server v.2.0 (Krogh et al., 2001); (e) the presence of N-terminal signal peptides with SignalP Server v.5.0 (Armenteros et al., 2019); and (f) gene ontology (GO) term IDs using Blast2GO software (Conesa & Götz, 2008) with the SwissProt Viridiplantae protein dataset (Consortium, 2019).

### 2.4. Duplication events in the rubber tree kinome

We determined duplication events of the PK superfamily in Hbr based on the physical location of PK genes and their compositional similarities assessed through comparative alignments with the Hbr genome using the BLASTn algorithm (Altschul et al., 1990). Tandem duplications were defined as PK pairs separated by a maximum distance of 25 kb on the same chromosome and with the following: (i) a minimum similarity identity of 95%; (ii) an E-value cutoff of 1.0E-10; and (iii) a 75% minimum sequence length coverage. The chromosomal location of putative tandemly duplicated PK genes was illustrated using MapChart v.2.32 (Voorrips, 2002), and synteny relationships of the PKs were visualized using Circos software v.0.69 (Krzywinski et al., 2009).

### 2.5. Transposable element search

We searched for transposable elements (TEs) in the Hbr genome using TE data of 40 plant species obtained from the PlanNC-TE v3.8 database (Pedro et al., 2018). For this purpose, we performed a comparative alignment between the TEs retrieved and the *H. brasiliensis* reference chromosomes using BLASTn (Altschul et al., 1990) for short sequences (blastn-short) with the following parameters: (i) minimum coverage of 75%; (ii) word size of 7; and (iii) an E-value cutoff of 1.0E-10. We selected TEs located within a 100 kb window from Hbr PK genes. The chromosomal localization of TEs was illustrated using MapChart v.2.32 (Voorrips, 2002).

### 2.6. RNA-Seq data collection

Several publicly available Hbr RNA-Seq experiments were collected from the NCBI Sequence Read Archive (SRA) database (Leinonen et al., 2010). The samples consisted of a wide range of tissues and comprised various genotypes. In total, we obtained 129 samples from 10 studies (Cheng et al., 2018; Deng et al., 2018; Lau et al., 2016; Li et al., 2016; Mantello et al., 2019; Montoro et al., 2018; Rahman et al., 2019; Sathik et al., 2018; Tan et al., 2017; Tang et al., 2016) that evaluated control and stress conditions (cold, drought, latex overexploitation, jasmonate, and ethylene treatments).

### 2.7. Expression analysis

The raw sequence data were submitted to a sequence quality control assessment using the FastQC tool (Andrews, 2010), following a low-quality read filtering and adapter removal step using Trimmomatic software v.0.39 (Bolger et al., 2014). After removing adapter sequences, we retained only reads larger than 30 bp and bases with Phred scores above 20. The corresponding coding sequences (CDSs) from Hb chr and Hb scaf were used as a reference for the quantification step using Salmon software v.1.1.0 (Patro et al., 2017) with the k-mer length parameter set to 31. The expression values of each PK transcript were normalized using the transcript per million (TPM) metric, and samples containing biological replicates were combined by defining the mean value among replicates. To visualize the expression of each kinase subfamily among different tissues and cultivars, we generated two heatmap figures for control and stressed samples using the R package pheatmap (Kolde & Kolde, 2015).

### 2.8. Coexpression network construction

Two coexpression networks of Hbr PK subfamilies, corresponding to control and abiotic stress situations, were modeled and visualized using the R package igraph (Csardi et al., 2006) with the minimum Pearson correlation coefficient set to 0.7. To assess the structure of each network and specific subfamily attributes, we estimated the hub scores of each PK subfamily from Kleinberg’s hub centrality scores (Kleinberg, 1999) and edge betweenness values from the number of geodesics passing through each edge (Brandes, 2001).

## 3. Results

### 3.1. Genome-wide identification, classification and characterization of PKs

Based on the established pipeline, we identified 2,842 typical putative PK genes in Hbr (Supplementary Table S1a), 1,531 in Mes (Supplementary Table S1b), and 863 in Rco (Supplementary Table S1c). The rubber tree kinome resulted from 1,206 (42.43%) proteins from the Hb scaf dataset and 1,636 (57.57%) from Hb chr. Interestingly, we also identified several PKs containing multiple kinase domains in all three datasets (191, 91, and 44 in Hbr, Mes and Rco, respectively) (Supplementary Tables S2), which were probably retained during evolution due to their action with specific substrates. Typical PKs were defined as protein sequences presenting high similarity to a given kinase domain with minimum coverage of 50%. The atypical PKs of Hbr (728), Mes (230), and Rco (95) were removed from subsequent analyses, being considered as probable pseudogenes.

Typical Hbr, Mes, and Rco PKs were further classified into groups and subfamilies based on the HMM profiles of 127 kinase subfamilies defined by Lehti-Shiu & Shiu (2012). The PK domain classification was validated by phylogenetic analyses (Supplementary Figs. S1-S2). Thus, PKs were grouped into 20 major groups: PKs A, G and C (AGC), Aurora (Aur), budding uninhibited by benzimidazoles (BUB), calcium- and calmodulin-regulated kinases (CAMK), casein kinase 1 (CK1), cyclin-dependent, mitogen-activated, glycogen synthase, and CDC-like kinases (CMGC), plant-specific, inositol-requiring enzyme 1 (IRE1), NF-kB-activating kinase (NAK), NIMA-related kinase (NEK), Pancreatic eIF-2*α* kinase (PEK), Receptor-like kinase (RLK)-Pelle, *Saccharomyces cerevisiae* Scy1 kinase (SCY1), Serine/threonine kinase (STE), Tyrosine kinase-like kinase (TKL), Tousled-like kinases (TLK), Threonine/tyrosine kinase (TTK), Unc-51-like kinase (ULK), Wee1, Wee2, and Myt1 kinases (WEE), and with no lysine-K (WNK). We also identified 72 PKs in Hbr (2.5%), 30 in Mes (2.0%), and 22 in Rco (2.5%) that did not cluster in accordance with any subfamily classification and were placed in the “Unknown” category (Supplementary Tables S3). This category represents species-specific PKs, which might be related to new gene families.

The RLK-Pelle was the most highly represented group in all three species, as evidenced in Fig. 1, and was divided into 59 different subfamilies, accounting for 65.5%, 68.1%, 65.2% of all rubber tree, cassava, and castor plant PKs, respectively, followed by the CMGC (6.4% in Hbr, 5.9% in Mes, 7.5% in Rco), CAMK (5.9% in Hbr, 6.5% in Mes, 6.5% in Rco), TKL (4.9% in Hbr, 4.9% in Mes, 5.6% in Rco) and others (Supplementary Table S4). Such similarity in the distribution of PK subfamilies corroborates the phylogenetic proximity between these species and the high conservation of PKs. We investigated the chromosomal positions, intron distribution and structural properties of Hbr, Mes, and Rco PKs using several approaches (Supplementary Tables S5-S7). Hbr and Mes PK genes were distributed along all Hbr and Mes chromosomes (Supplementary Fig. S3) with an apparent proportion to the chromosome length. There was also a higher concentration of PKs in the subtelomeric regions, which may indicate chromosomal rearrangements and increased variability. Most PK genes contained at least 1 intron, and only 284 (10.0%), 229 (14.9%), and 140 (16.2%) intronless genes were found in Hbr, Mes, and Rco, respectively. In specific subfamilies, the distribution of introns presented a similar profile, e.g. the CK1 CK1 subfamily (21 members) had an average of 14 introns per gene with a coefficient of variation of about 10%. Interestingly, members of the SCY1 SCYL2, TKL-Pl-7, RLK-Pelle LRR-XIIIb, and RLK-Pelle LRR-VII-3 subfamilies presented the same quantity of introns, which might be related to conserved plant roles across evolution.

**Fig. 1.**
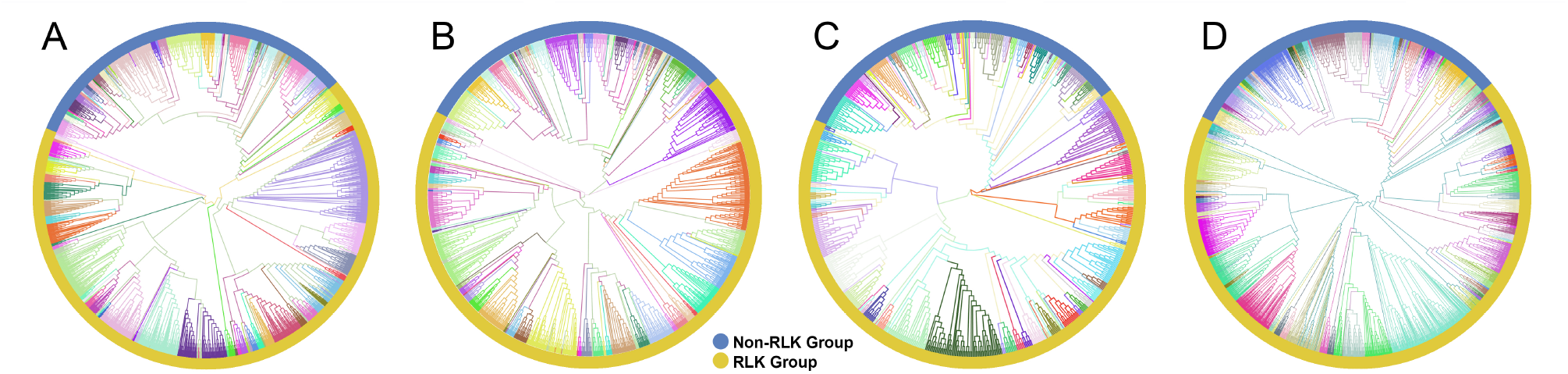
Phylogenetic analyses of putative typical protein kinases (PKs) identified in the *Hevea brasiliensis* (Hbr), *Manihot esculenta* (Mes), and *Ricinus communis* (Rco) genomes. (A) Phylogenetic tree constructed with 2,842 Hbr PKs organized into 123 subfamilies. (B) Phylogenetic tree of the 1,531 Mes PKs organized into 123 subfamilies. (C) Phylogenetic tree of the 863 Rco PKs organized into 125 subfamilies. (D) Phylogenetic tree of all Hbr, Mes, and Rco PKs. Kinase subfamilies are represented by different branch colors.

Most interestingly, the protein characteristics of all three kinomes were highly comparable. Several PKs had predicted transmembrane domains (45.6% in Hbr, 50.5% in Mes, and 48.2% in Rco) and N-terminal signal peptides (29.6%, 37.3%, and 33.5%, respectively). Similarly, the distribution of molecular weights and isoelectric points were relatively uniform (Supplementary Fig. S4), equally observed for the subfamily divisions. Moreover, the subcellular localization predictions performed with the selected software were mostly on the plasma membrane, cytoplasm and nucleus (Supplementary Fig. S5), in accordance with the enriched “cellular component” GO category (Supplementary Tables S8; Supplementary Fig. S5). Indeed, PKs have a more pronounced presence across the plasma membrane with specific members acting on other subcellular components.

Finally, we investigated the domain composition of PKs based on the complete set of conserved domains present in the Pfam database. In total, we identified 1,472 PKs containing additional conserved domains in Hbr (52.8%), 827 in Mes (54.0%) and 442 in Rco (51.2%) (Supplementary Tables S9-S10). This varied profile confirms the diverse functions of PKs, their speciation and importance for plant adaptation. Interestingly, we identified a significant portion of members from groups CAMK (58.0% in Hbr, 61.6% in Mes, and 64.3% in Rco), RLK-Pelle (63%, 65.5%, and 63.9%, respectively), and TKL (48.2%, 48%, and 52%) with additional domains. Based on these findings, it can be observed that the evolutionary history of PKs is also based on specific domain arrangements.

### 3.2. Kinase duplication events in H. brasiliensis

To examine the expansion of PK subfamilies, tandemly duplicated kinase genes were identified based on their physical localization on Hbr chromosomes, and protein similarities were assessed through comparative alignments. Taken together, we found that 339 of the 2,842 Hbr PK genes (~11.9%) were arranged in clusters of highly similar gene sequences among the 18 reference chromosomes, which are likely to represent tandem duplication events (TDEs) of the kinase superfamily in rubber tree (Fig. 2A). These genes were dispersed in 145 separate clusters and comprised members of 63 kinase subfamilies (Supplementary Tables S11-S12). Chromosome 14 showed the highest number of TDEs (19), containing 47 PK genes. In contrast, chromosome 1 contained the least number of TDEs (2). We found that for some kinase subfamilies, a large portion of their members originated from TDEs, suggesting that such duplication events influenced the expansion of specific groups due to putative functional implications. A total of 100%, 100%, and 75% of TTK, ULK Fused, and CMGC CDKL-Os members were tandemly organized, while other subfamilies, such as RLK-Pelle DLSV, showed the largest absolute number of TDEs (45) distributed across 9 chromosomes, although it accounted for only 16.8% (45/268) of its total size.

**Fig. 2.**
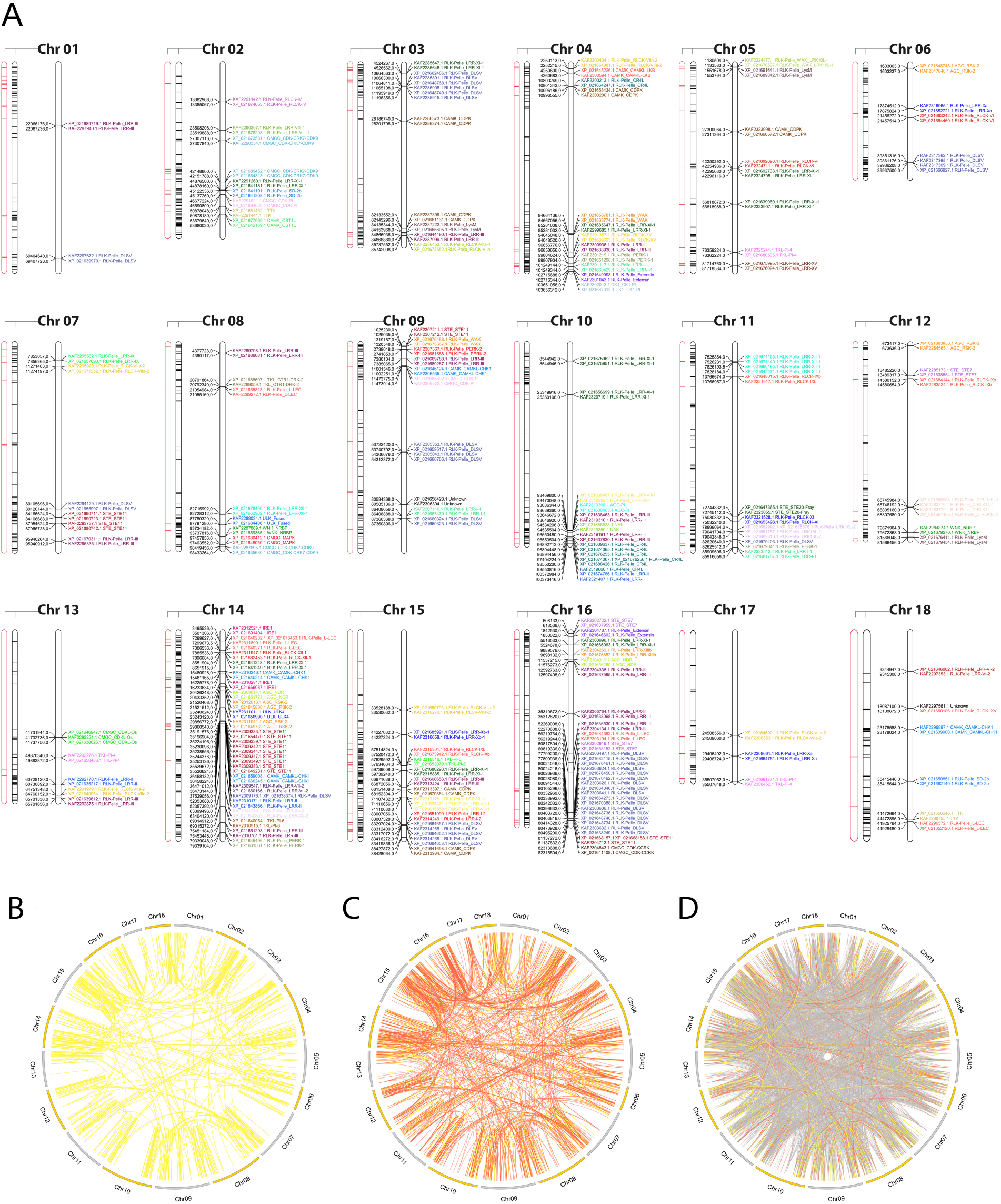
(A) Kinase distribution along *Hevea brasiliensis* chromosomes. For each chromosome, from left to right: (i)transposable elements located within a 100 kb window around kinase genes are highlighted in red; (ii) all genes with kinase domains are highlighted in black; and (iii) tandemly duplicated kinase genes are colored and labeled according to the kinase subfamily classification. (B) Potential segmental duplication events in the *H. brasiliensis* genome considering similarities greater than 90% (yellow); (C) 75% (orange); and (D) 50% (gray).

Segmental duplication events were estimated based on sequence similarities between two or more PKs separated by a genomic window larger than 100 kb or present in different chromosomes. Genomic correspondences increased as the sequence similarity decreased (Fig. 2D). In total, we identified 858 kinase correspondences with compositional similarity greater than 90%, 1,673 for 75% and 10,121 for 50%. Clearly, the expansion of the rubber tree kinome is mainly caused by segmental duplication events, probably related to the *Hevea* paleopolyploid genome origin (Pootakham et al., 2017). To further investigate potential biological processes associated with duplicated kinase genes, we performed a functional annotation pipeline on tandemly duplicated PKs and selected GO terms related to the “biological process” category (Supplementary Fig. S6). The findings were consistent with those resulting from the analysis performed using the complete set of Hbr PKs (Supplementary Fig. S7). Although there is a subset of PK subfamilies with members originated by TDEs, their functional profile represents the entire repertoire of PK GO terms, which shows that, even less pronounced, TDEs in rubber tree play an important role in the kinome expansion.

### 3.3. Transposable elements in H. brasiliensis genome

Due to the high abundance of TEs in the rubber tree genome and current evidence associating these elements with phenotypic modifications (Francisco et al., 2021; Wu et al., 2020), we predicted TEs near Hbr PK genes using a comprehensive database that combined data from overlapping regions of TE features and several classes of noncoding RNAs (ncRNAs). Overall, the percentage of TEs associated with PK genes in the rubber tree was reduced (23.7%) when compared to overlapping ncRNAs (76.3%) (Supplementary Table S13), which have a broader set of regulatory roles (Hadjiargyrou & Delihas, 2013). Out of the 8,457 annotated TEs in the reference genome, 88% were classified as long terminal repeat (LTR) retrotransposons. These elements appeared to be associated (within a 100 kb genomic window) with 362 (12.7%) kinase genes (Fig. 2A, Supplementary Table S14), of which 56 (15.5%) were tandemly duplicated. Such findings highlight the importance of these joint mechanisms for understanding the complex dynamics behind rubber tree kinome expansion. Nearly 73.2% of these duplicated genes were members of the RLK-Pelle group, reiterating its importance.

### 3.4. Expression patterns of PK subfamilies

In order to supply a broad evaluation of PK expression, we analyzed the expression levels of 118 kinase subfamilies among 129 samples related to control and different abiotic stress conditions (Supplementary Table S15). The resulting dataset comprised transcriptomic data of 14 different cultivars from various tissues and organs, including leaf, petiole, bark, latex, seed, male and female flowers, in early and mature developmental stages. After filtering out low-quality reads and removing adapter sequences, we mapped the filtered reads to the complete set of CDS sequences in Hb chr and Hb scaf reference genomes separately and further generated a subset of the quantifications corresponding to PK genes present in the Hbr kinome. Such an approach enabled a deep characterization of PK gene expression, yet under-explored. The results were normalized to TPM values (Supplementary Tables S16-S17), and 3 samples presenting significantly low quantifications were excluded (Hb_Bark_3001_normal_rep1, Hb_Latex_712_normal_rep2, Hb _Latex_2025_normal_rep1). For cases where replicates were present, the expression values were averaged.

For both control and stress heatmaps (Supplementary Figs. S8-S9, respectively), samples belonging to the same tissues were clustered together based on Euclidean distance measures. In general, we observed similar patterns of expression of each kinase subfamily within samples of a given tissue; however, specific experimental conditions of each RNA-Seq dataset may have influenced the expression levels, leading to inconsistent patterns in some cases. In addition to the high genetic diversity among the rubber tree genotypes used in the datasets, some PK subfamilies may have a different response to each type of stress present in our study, as observed in the different clustering profiles (Supplementary Fig. S10). From left to right in the heatmap containing all experiments (Supplementary Fig. S10), there were 5 major clusters separated into the following categories: (i) latex; (ii) leaf and seed tissues; (iii) bark, root, male and female flowers; (iv) leaf; and (v) samples from latex and petiole.

Interestingly, several subfamilies were highly expressed in nearly all samples, including AGC PKA-PKG, TKL-Pl-1, RLK-Pelle LRR-VIII-1, RLK-Pelle RLCK-IXa, and Aur, which probably represent PK subfamilies not affected by stress stimuli, but related to basal plant functions. Therefore, distinctions in PK expression between leaf and latex samples were clear. Latex and bark tissues presented lower expression in most subfamilies; however, we detected a few cases where the expression in latex and bark was significantly higher than in leaves, such as STE STE-Pl and RLK-Pelle LRR-VIII-1, providing indicative targets of the roles of specific PK subfamilies that affect the secondary metabolism. Additionally, we found a small number of subfamilies with elevated expression in bark (TKL-Pl-7 and ULK ULK4), which depends directly on the seasonality of the climate (Budzinski et al., 2016). Overall, in the analysis of the expression levels under abiotic stress conditions, the number of subfamilies that presented moderately high (dark orange) and high (red) expression increased when compared to control samples (highlighted in blue), showing the activation of PK subfamilies under stress.

### 3.5. Coexpression networks in response to abiotic stresses

The quantification analysis revealed different expression profiles of PK subfamilies among different tissues, genotypes and conditions. To expand our understanding of how these proteins interact under exposure to abiotic conditions, we further investigated potential relationships between kinase subfamilies by constructing coexpression networks based on the expression data described above. Using the Hbr PK set, two independent networks were constructed: one for control and one for abiotic stress conditions. Using such an approach, we were able to investigate PK subfamily interactions activated under abiotic stress conditions. For each network, we used the following conventions: (i) kinase subfamilies were represented by separate nodes; the node size corresponded to the mean gene expression value; (iii) the edges represented coexpression events determined by pairwise expression correlations between subfamilies with a minimum Pearson correlation coefficient of 0.7; and (iv) the edge thickness corresponded to the degree of correlation, from moderate (minimum PCC of 0.7) to relatively strong (minimum PCC of 0.9) correlations.

We observed a different number of edges between networks (1,162 in control and 704 in stress). The presence of a reduced number of connections in the stress coexpression network might indicate the impact of stress conditions on PK subfamilies, changing their expression profile and consecutively their functional interaction with other subfamilies. Moreover, we found 15 elements in each network that were disconnected from the main core (i.e., kinase subfamilies with no significant correlation in the expression); however, they were related to different subfamilies in each network (Fig. 3), showing that under specific conditions PK subfamilies might activate a different communication mechanism, overcoming external factors. Figs. 3B and 3D highlight the red correlation similarities between control and stress coexpression networks, while dark grey edges represent unique connections for each condition. It is possible to observe that there is a common core of connections between the two modeled networks, with specific interactions related to the samples used to model these systems, i.e. the presence or absence of stress.

**Fig. 3.**
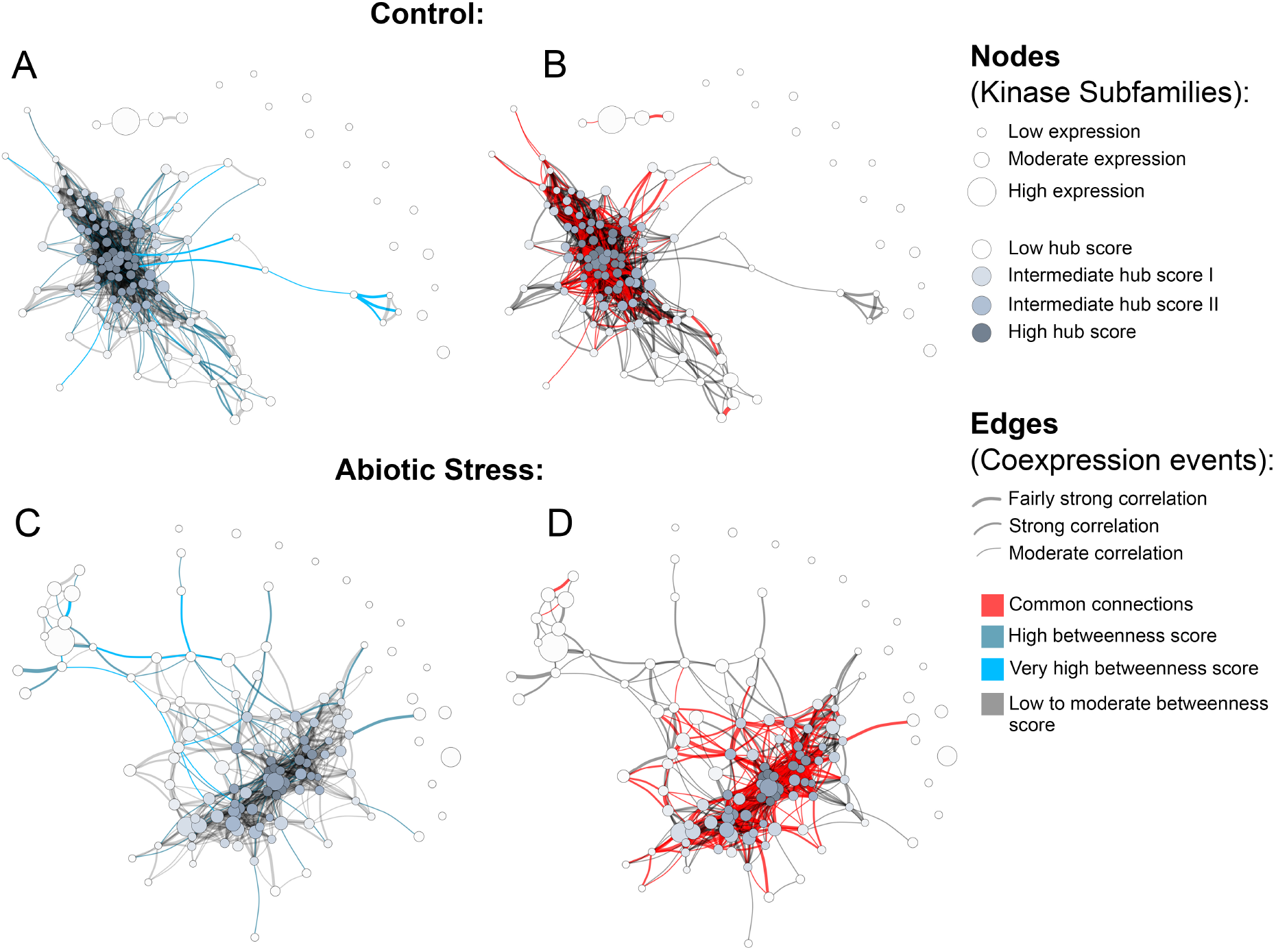
Coexpression networks for *H. brasiliensis* (Hbr) kinase subfamilies. (A) Hbr control network with betweenness values highlighted in light blue. (B) Hbr control network indicating edge similarities (red) with the Hbr stress network. (C) Hbr abiotic-stress network with betweenness values highlighted in light blue. (D) Hbr abiotic stress network indicating edge similarities (red) with the Hbr control network.

To obtain an overview of the most influential subfamilies in PK processes, we first calculated hub centrality scores within each network, which are represented by the node colors in Fig. 3 (Supplementary Tables S18). Elevated hub scores (highlighted in dark gray in Fig. 3) indicate PK subfamilies with a significant number of connections, probably developing a broad set of molecular interactions and being an important component in PK communication. Interestingly, out of the 10 subfamilies with the highest hub scores (ranging from 0.85 to 1) in the Hbr control network, eight belonged to members of the RLK-Pelle group (RLK-Pelle L-LEC, RLK-Pelle RLCK-V, RLK-Pelle RLCK-VIIa-2, RLK-Pelle LRR-I-1, RLK-Pelle LysM, RLK-Pelle SD-2b, RLK-Pelle DLSV, and RLK-Pelle RLCK-VIIa-1), while the others were CAMK-CDPK and CK1 CK1-Pl. In addition to being the most abundant and associated with TDEs, the importance of the RLK-Pelle group is evidenced by its putative interaction with several PK subfamilies. In contrast, under adverse conditions, the hub scores of the top 10 sub-families varied from 0.75 to 1; only 6 of them were members of the RLK-Pelle group (RLK-Pelle RLCK-X, RLK-Pelle PERK-1, RLK-Pelle LysM, RLK-Pelle SD-2b, RLK-Pelle L-LEC, RLK-Pelle RLCK-VIIa-1), while the others were CAMK CDPK, TKL-Pl-4, STE STE7, and CK1 CK1-Pl.

Ultimately, we investigated network structural weaknesses by measuring edge betweenness centrality scores among kinase subfamily interactions (edges) within each network (Supplementary Tables S19). The edges presenting elevated betweenness values (colored in light blue in Fig. 3) indicate relationships sustained by few connections possibly related to a greater flow of interaction into the complex system modeled. Such connections represent specific interactions between PK subfamilies with a significant importance for the whole system communication, i.e. subfamilies acting in different biological processes through indirect implications. Overall, under adverse conditions, PK subfamilies tended to arrange into a less cohesive network architecture, as evidenced by a large number of scattered connections. Through betweenness measures, we observed that other influential subfamily pairs of the control network were RLK-Pelle LRR-XII-1/RLK-Pelle RLCK-XVI, RLK-Pelle LRR-VII-3/RLK-Pelle RLCK-XVI, and RLK-Pelle CR4L/RLK-Pelle LRR-Xb-1, in contrast to the ones found during abiotic stress situations: RLK-Pelle RLCK-Os/TKL-Pl-6, CAMK CAMK1-DCAMKL/RLK-Pelle RLCK-Os, and RLK-Pelle RLCK-V/TKL-Pl-6. These differences in the top betweenness connections demonstrate the distinct interactions of PK subfamilies during the stress response.

## 4. Discussion

### 4.1. Rubber Tree Kinome

In the last decades, an increasing number of initiatives have been established to produce a high-quality reference genome for the rubber tree (Liu et al., 2020b; Pootakham et al., 2017; Rahman et al., 2013; Tang et al., 2016). However, the high complexity of the Hbr genome introduces many challenges that have hampered the ability to obtain contiguous genomic sequences and complete gene annotation (English et al., 2012; Pootakham et al., 2017). Although the most recent version of the Hbr genome (Liu et al., 2020b) provided the assembly of contiguous sequences for chromosomes for the first time, there is still a lack of knowledge about its gene content and functional implications, highlighting the need for efforts to profile and fully characterize important protein families, such as PKs. Here, we established a combined approach to generate a comprehensive and diverse kinase database for Hbr. Joining two independent rubber tree genomic resources (Liu et al., 2020b; Tang et al., 2016) and comparing them with kinomes from two other members of the *Euphorbiaceae* family (Mes and Rco) enabled an in-depth investigation of the rubber tree kinome, supplying a large and reliable reservoir of data. We suggest that such a combination approach could be a valuable strategy for other species with limited genomic resources, increasing the identification and definition of important molecular elements. Plant kinomes have been studied in several other species, including 942 members in *A. thaliana* (Zulawski et al., 2014), 2,166 in soybean (Liu et al., 2015), 1,168 in grapevine (Zhang, 2003), and 1,210 in sorghum (Aono et al., 2021). In this study, we identified 2,842 PKs in the rubber tree, a considerably larger size when compared to 1,531 in cassava and 863 in castor bean. Although the rubber tree possesses a large genome (1.47 Gb), which is nearly 3-4.5 times larger than the cassava (495 Mb) and castor bean (320 Mb) genomes (Bredeson et al., 2016; Chan et al., 2010; Tang et al., 2016), the large kinome size in Hbr resulted from the combination of two sources of PKs related to different rubber tree genotypes (Reyan7-33-97 and GT1) (Liu et al., 2020b; Tang et al., 2016). When analyzing the two Hbr PK sets separately, we found a much smaller number of PKs in each of them (1,809 and 1,379), placing the Hbr kinome in the range of other plant species. The discrepancy found between the two sources of data reinforces the differences in completeness among genome assemblies, which we believe that could potentially mislead further genomic investigations, especially considering the elevated heterozygosity levels and the high amount of repetitive elements in the rubber tree genome (Gouvêa et al., 2010; Lau et al., 2016), mainly caused by the demographic history and genetic diversity of the rubber tree. Additionally, a recent study showed that the number of PKs in sugarcane was significantly decreased on the allelic level when compared to those found for all allele copies of a given gene (Aono et al., 2021), demonstrating that the redundancy in PK datasets may contribute to the overestimation of kinome sizes.

Similar to the results of other kinome studies (Aono et al., 2021; Ferreira-Neto et al., 2021; Liu et al., 2020a, 2015; Wei et al., 2014; Yan et al., 2018; Zhu et al., 2018a; Zulawski et al., 2014), RLK-Pelle was the most pronounced group in Hbr, Mes and Rco kinomes. Considering the diverse functions of PKs (Lehti-Shiu & Shiu, 2012), our study corroborates the remarkable role of this group in the Hbr response to stress. We also suggest that such importance transcends its high abundance, including its expansion through TDEs and also the way the RLK-Pelle subfamilies functionally interact with other PKs, as evidenced by the coexpression relationships found. One notable feature observed across Hbr PKs was the large diversity in domain configuration. Most PKs (56.9%) in rubber trees had two or more functional domains incorporated with them, similar to what has been observed in cassava (59.2%), castor bean (55%), soybean (56.5%) (Liu et al., 2015), and pineapple (50.7%) (Zhu et al., 2018a). By combining different protein properties, we inferred a considerable presence of extracellular domains (ECDs), mainly because of: (i) the large diversity of additional domains in PK genes; (ii) the detection of signal peptides and transmembrane regions; and (iii) the wide range of subcellular localizations predicted (Supplementary Fig. S5). PKs combined with ECDs may broaden the scope of functionality within signaling networks by sensing new extracellular signals and their aggregation to existing response networks (Gish & Clark, 2011; Lehti-Shiu & Shiu, 2012), which makes our results a valuable source of data for establishing PKs sensing different environmental stimuli and signalling important metabolic mechanisms, such as the rubber biosynthesis.

Our comparative analyses of the PKs of the three *Euphorbiaceae* species revealed a high degree of similarity in their kinase subfamily compositions, protein characteristics and gene organization (Supplementary Figs. S2-S4). Our integrative approach allowed us to corroborate the validity of the Hbr kinome, which was composed of different data sources and had evident resemblances with closely related phylogenetic species. However, Mes was more similar to Hbr in kinome size than Rco. This pattern of gene expansion within the *Euphorbiaceae* clade was also observed in other gene families, including the SWEET and SBP-box families (Cao et al., 2019; Li et al., 2019). Although the focus of our study was not to evaluate the PK evolutionary divergences between Hbr, Mes and Rco, the similarities that we observed between the kinomes are corroborated by the evolutionary history of such taxonomic groups. Phylogenetic studies indicated that *Hevea* and *Manihot* underwent a whole-genome duplication event before their divergence of approximately 36 million years ago (MYA), while the *Ricinus* lineage diverged from other Euphorbia members approximately 60 MYA (Bredeson et al., 2016; Shearman et al., 2020). In this sense, the increase in the size of the PK superfamily could be partially attributed to the expansion of several gene families through duplication events during the evolutionary history of these species, as already reported by Lehti-Shiu & Shiu (2012).

### 4.2. Duplication Events in the Rubber Tree Kinome

Our analysis suggested that segmental duplications mostly accounted for Hbr kinome expansion (Fig. 2B), with ~58.9% of PKs displaying more than 75% compositional similarities. TDEs, on the other hand, seemed to contribute to the expansion of the PK superfamily to a lesser extent and were restricted to a few subfamilies (Supplementary Table S11), accounting for the generation of nearly 11.9% of PK genes in Hbr. As we observed the same functional profile in the subgroup of PKs tandemly duplicated and throughout the kinome, we infer that such TDEs might be important for the maintenance of PK activities throughout evolution. This TDE rate was within the range of what has been reported for other higher plants, such as 10.6% in soybean (Liu et al., 2015), 12.5% in pineapple (Zhu et al., 2018a), and 14.8% in strawberry (Liu et al., 2020a).

Tandemly duplicated kinases have been associated with stress responses (Freeling, 2009), which is of great interest for molecular breeding. Taking this association into account, we suggest an important role of RLK-Pelle DLSV, RLK-Pelle LRR-III, RLK-Pelle LRR-XI-1, STE STE11, and CAMK CDPK subfamilies in stress signaling, which should be investigated in further studies. Additionally, in rubber tree research, different initiatives have brought to light the importance of PKs in the configuration and maintenance of economically important traits, including not only resistance to different types of stress (Duan et al., 2010; Jin et al., 2017; Mantello et al., 2019; Venkatachalam et al., 2010) but also plant performance in the field (Bini et al., 2022; Francisco et al., 2021). Together with these findings, recent contributions have pinpointed the role of TEs beyond Hbr genomic organization, suggesting a potential influence on the configuration of desirable rubber tree traits (Francisco et al., 2021; Wu et al., 2020). We also identified a considerable number of PKs (15.5%) associated with TEs, which we also suggest is related to the kinome expansion. In Hbr, Wu et al. (2020) showed that TEs located in gene regulatory regions were involved in latex production through cis regulation, which would explain the differential gene expression among contrasting genotypes. The incidence of PKs close to TEs pinpoints the importance of such elements on PK functionality, as already demonstrated by other studies describing TE-mediated regulation in kinases (Fan et al., 2019; Zayed et al., 2007).

It has been well established that TEs are abundant in the rubber tree genome, and the proportion of TE types found in our study was similar to those found by other authors (Liu et al., 2020b; Tang et al., 2016; Wu et al., 2020). Due to the mutagenic potential of TEs caused by epigenetic mechanisms, these elements can alter regulatory networks and confer genetic adaptations, leading to important phenotypic variations (Lisch, 2013; Wei & Cao, 2016; Wu et al., 2020); this has currently received great attention in genetic improvement programs for several species (Domínguez et al., 2020; Lee et al., 2006; Wang et al., 2020). Similar to PKs, which are especially active during abiotic stress (Jaggi, 2018; Morris, 2001), TEs are related to plant adaptations throughout evolution (Casacuberta & González, 2013; Dubin et al., 2018; Lisch, 2013; Naito et al., 2009; Negi et al., 2016). We found an association between TEs and specific kinase subfamilies, such as those present in the RLK-Pelle group. As expected due to the occurrence of duplication events caused by TEs (Flagel & Wendel, 2009), we also observed an association of these elements with tandemly duplicated PKs, enabling the elucidation of diverse biological mechanisms favoring stress resistance.

Considering the growing demand for latex production, the interest in developing cold-resistant cultivars for expanding *Hevea* plantation motivates the comprehension of the molecular mechanisms associated with resistance. Such a scientific challenge is directly related to the signaling mechanisms of PKs, with the direct action of specific subfamilies. We believe that the association of TEs, TDEs and specific PK subfamilies provides deeper insights into the role and maintenance of PKs for stress resilience. In addition to the genetic adaptation of the rubber tree, TEs have increasingly been attracting attention, mainly due to their association with (i) intron gain (Gozashti et al., 2022); (ii) increased heterozygosity in stress-responsive genes (De Kort et al., 2022); (iii) increases in gene expression variation (Uzunović et al., 2019); and (iv) important polymorphisms associated with the genetic effects of complex traits(Vourlaki et al., 2022). These facts, together with the association of TDEs and stress responses, enabled the establishment of important PK subfamilies with high potential to be investigated for climate resilience.

### 4.3. Gene Expression Evaluations

Differentially expressed gene (DEG) analyses are based on statistical tests performed on gene expression quantifications measured under certain conditions, contrasting physiological contexts and different stimuli, enabling the evaluation of increased gene expression (Casassola et al., 2013; Costa-Silva et al., 2017). Although we did not perform such an analysis of Hbr PKs due to the different experiments and datasets employed, it was possible to visualize a distinct overall expression profile for subfamily expression across the samples employed, illustrating putative molecular mechanisms adopted by PK subfamilies to overcome stress conditions (Mantello et al., 2019).

The subfamilies CAMP AMPK, CMGC PITthe, CK1 CK1, RLK-Pelle LRR-XIIIb, and RLK-Pelle URK-2 exhibited more pronounced expression in samples under stress conditions, as already reported in other studies (Hawley et al., 2005; Saito et al., 2019). Even without statistical evaluations, it is possible to infer an impact of stress conditions on the molecular mechanisms of these PK subfamilies. CAMP AMPK has been described as an important energy regulator in eukaryotes, coordinating metabolic activities in the cytosol with those in mitochondria and plastids, possibly allocating energy expenditure to overcome these adversities (Hawley et al., 2005; Roustan et al., 2016; Suzuki et al., 2012). Although members of the CK1 subfamily were highly conserved in eukaryotes and involved in various cellular, physiological and developmental processes, their functions in plant species are still poorly understood (Saito et al., 2019). Studies indicated that CK1 members in *A. thaliana* are involved in several processes related to the response to environmental stimuli, such as regulation of stomatal opening (Zhao et al., 2016), signaling in response to blue light (Tan et al., 2013), organization and dynamics of cortical microtubules (Ben-Nissan et al., 2008), and ethylene production (Tan & Xue, 2014). In this context, our study corroborates the putative role of the CK1 subfamily, highlighting its change in expression during stress response.

As a complementary approach to elucidate different patterns in the expression of PK subfamilies in control and abiotic stress-related samples, we employed gene coexpression networks. Through a graph representation of the PK subfamily interactions in these two groups of samples, we estimated coexpression patterns inherent to each network, inferring functional implications through the network topology. As the modeled coexpression networks represent the interaction between the PK subfamilies, we suggest that, by contrasting the topological differences between these networks, it is possible to infer changes in the interaction of the PK subfamilies induced by stress conditions. Even with a common set of interactions (Fig. 3B and 3D), it is possible to note fewer associations in the network modeled with stress samples (a loss of

~40%). During stress conditions, PKs act through a signaling system to activate specific cellular responses to overcome these adversities, affecting molecular interactions. In the network modeled with samples affected by stress, it is possible to visualize such alterations, i.e. the cohesive set of interactions between the PK subfamilies becomes more sparse, evidencing that specific subfamilies start to perform different functions.

In a complex network structure, nodes with the largest number of connections (high degree) are called hubs, which are elements recognized as critical to network maintenance (Barabasi & Oltvai, 2004). Therefore, PK subfamilies with the highest hub scores are considered to be important regulators over the set of biological mechanisms affected by PKs (Azuaje, 2014; Barabasi & Oltvai, 2004; Van Dam et al., 2018), which provides additional insights into key mechanisms over PKs’ action (Vandereyken et al., 2018).

In both networks modeled, we found that the CAMK CDPK subfamily had the highest hub score, suggesting the importance of calcium signals over Hbr PK activities, as already reported in soybean (Liu et al., 2015). Interestingly, members of the CAMK CDPK subfamily were found to be tandemly duplicated and also associated with TEs. Considering the previously described importance of TDEs and TEs in the Hbr kinome, we suggest that such a subfamily plays a key role in activating other PK subfamilies to overcome the stress response. Members of the RLK-Pelle group were also identified as hubs in both networks, reinforcing the primary and secondary metabolic functions of this group (Bolhassani et al., 2021). Additionally, TKL-Pl-4 was among the hubs in the stress-related network, corroborating the already described upregulation of members of this subfamily in stress conditions (Yan et al., 2017). Similar to CAMK CDPK, the TKL-Pl-4 subfamily was also related to TDEs and TEs, which shows the conservation of such a subfamily throughout evolution. Furthermore, under stress conditions, we observed that the TKL-Pl-4 subfamily starts to play a more prominent role in the network, probably indicating its signaling activity on other PK subfamilies in response to stress.

Another measure evaluated in the modeled networks was edge betweenness scores. In a complex network structure, edges with high betweenness indicate points of vulnerability in the network structure, i.e. connections that, if removed, have a larger probability of causing network separation. We suggest that in the networks modeled for PK subfamily interactions, the identification of such edges can supply indicators of subfamilies mediating a significant amount of mechanisms over a larger set of PKs. Additionally, the comparison of edges with high betweenness between networks can provide clues about the change of molecular mechanisms affected by stress. Interestingly, the two highest betweenness values for the control network (RLK-Pelle LRR-XII-1/RLK-Pelle RLCK-XVI and RLK-Pelle LRR-VII-3/RLK-Pelle RLCK-XVI edges) were disrupted in the stress-associated network. The RLK-Pelle RLCK-XVI sub-family did not have any connection in the stress network, showing a change in the interaction of this subfamily under stress. We infer that such a change is caused by stress conditions, i.e. the putative functions of the RLK-Pelle RLCK-XVI subfamily are related to biological processes that are directly affected by the external stimuli. Interestingly, RLCK members have already been shown to be related to plant growth and vegetative development (Gao & Xue, 2012; Yan et al., 2018), which is directly impacted by stress. We suggest that such a subfamily is an interesting target for understanding the impact of climate changes on Hbr.

Additionally, in the stress-related network, we found that the PK subfamily pairs RLK-Pelle RLCK-Os — TKL-Pl-6, CAMK CAMK1-DCAMKL — RLK-Pelle_RLCK-Os, and RLK-Pelle_RLCK-V — TKL-Pl-6 had the largest betweenness scores. All these subfamilies presented a similar number of connections in the control network; however, they were not considered vulnerability points. This result indicated that under stress, established connections might become sensitive and cause network breaks in more adverse conditions, representing PK subfamilies that can have their functions altered under high stress, such as the RLK-Pelle RLCK-XVI subfamily in the change from control to stress. Given the described downregulation of TKL-Pl-6 expression during stress (Yan et al., 2017), our results corroborate the impact of stress not only on its activity, but also on the way this subfamily interacts with other PK subfamilies. Additionally, CAMK CAMK1-DCAMKL has already been described as induced during stress (Liu et al., 2015), and such a fact can be observed in our network, with changes in the network configuration.

Given the importance of rubber trees, the rising demand for latex production, and the elevated complexity of the Hbr genome (Board, 2018; Tang et al., 2016), providing resources for understanding stress responses is of great interest for Hbr breeding programs (Priyadarshan & Goncalves, 2003). Our work provided a rich and large reservoir of data for Hbr research. In the first study to profile the complete set of PKs in Hbr, we combined different data sources to provide a wider PK characterization, taking advantage of the resources available and contrasting our results with two phylogenetically close species. From a set of 2,842 PKs classified into 20 groups and distributed along all Hbr chromosomes; our findings demonstrated the high diversity and scope of functionality of Hbr PKs. Additionally, we provided different insights across stress responses in rubber trees through the association of tandemly duplicated PKs, TEs, gene expression patterns, and coexpression events.

## Supporting information

Supplementary Results

## Funding

This work was supported by grants from the Fundação de Amparo á Pesquisa do Estado de São Paulo (FAPESP), the Conselho Nacional de Desenvolvimento Científico e Tecnológico (CNPq) and the Coordenação de Aperfeiçoamento de Pessoal de Nível Superior (CAPES, Computational Biology Programme). LBS received an undergraduate fellowship from FAPESP (2019/19340-2); AA received a PhD fellowship from FAPESP (2019/03232-6); FF received a PhD fellowship from FAPESP (2018/18985-7); and AS received research fellowships from CNPq (312777/2018–3).

## Acknowledgments

We would like to acknowledge the Fundação de Amparo à Pesquisa do Estado de São Paulo (FAPESP), the Conselho Nacional de Desenvolvimento Científico e Tecnológico (CNPq), and the Coordenação de Aperfeiçoamento de Pessoal de Nível Superior (CAPES).

## Author contributions

LBS, AA and FF performed all analyses and wrote the manuscript. AS, CS and LMS conceived of the project. All authors reviewed, read and approved the manuscript.

## Data Availability Statement

The datasets analyzed for this study can be found in the NCBI Sequence Read Archive (SRA) database (https://www.ncbi.nlm.nih.gov/bioproject), under project numbers: PR-JNA432826, PRJNA280256, PRJNA182078, PRJNA483203, PRJNA394848, PRJNA310171, PRJNA281775, PRJNA357682, PRJDB4387, and PRJNA511923.

## Supplementary Tables

**Table S1a**. Kinase domain annotation of 3,188 *H. brasiliensis* protein kinases.

**Table S1b**. Kinase domain annotation of 1,531 *M. esculenta* protein kinases.

**Table S1c**. Kinase domain annotation of 863 *R. communis* protein kinases.

**Table S2a**. *H. brasiliensis* kinase domain organization of 191 protein kinases containing multiple kinase domains.

**Table S2b**. *M. esculenta* kinase domain organization of 91 protein kinases containing multiple kinase domains.

**Table S2c**. *R. communis* kinase domain organization of 44 protein kinases containing multiple kinase domains.

**Table S3a**. Subfamily classification of *H. brasiliensis* protein kinases.

**Table S3b**. Subfamily classification of *M. esculenta* protein kinases.

**Table S3c**. Subfamily classification of *R. communis* protein kinases.

**Table S4**. The number of kinase genes in different subfamilies.

**Table S5a**. Characterization of *H. brasiliensis* protein kinases.

**Table S5b**. Characterization of *M. esculenta* protein kinases.

**Table S5c**. Characterization of *R. communis* protein kinases.

**Table S6a**. *H. brasiliensis* 2,842 kinase gene localizations and intron quantities.

**Table S6b**. *M. esculenta* 1,531 kinase gene localizations and intron quantities.

**Table S6c**. *R. communis* 863 kinase gene localizations and intron quantities.

**Table S7**. Chromosomal position estimation of 1,636 *H. brasiliensis* kinase genes.

**Table S8a**. Gene Ontology (GO) annotations for the 2,842 *H. brasiliensis* protein kinases.

**Table S8b**. Gene Ontology (GO) annotations for the 1,531 *M. esculenta* protein kinases.

**Table S8c**. Gene Ontology (GO) annotations for the 863 *R. communis* protein kinases.

**Table S9a**. Domain annotation of *H. brasiliensis* protein kinases.

**Table S9b**. Domain annotation of *M. esculenta* protein kinases.

**Table S9c**. Domain annotation of *R. communis* protein kinases.

**Table S10a**. Domain organization of *H. brasiliensis* protein kinases.

**Table S10b**. Domain organization of *M. esculenta* protein kinases.

**Table S10c**. Domain organization of *R. communis* protein kinases.

**Table S11**. Tandemly duplicated kinase genes across the *H. brasiliensis* genome.

**Table S12.** Pairs of tandemly duplicated kinases correspondences in the *H. brasiliensis* genome.

**Table S13**. Number of *H. brasiliensis* protein kinases near transposable elements.

**Table S14**. Description of *H. brasiliensis* kinase genes associated with transposable elements.

**Table S15**. *H. brasiliensis* RNA-Seq experiment organization (retrieved from the SRA database).

**Table S16**. *H. brasiliensis* kinase transcripts per million (TPM) values across samples.

**Table S17**. *H. brasiliensis* kinase quantifications across subfamilies.

**Table S18a**. Related information of the *H. brasiliensis* kinase subfamily control coexpression network.

**Table S18b**. Related information of the *H. brasiliensis* kinase subfamily stress coexpression network.

**Table S19a**. Edge betweenness values calculated across the *H. brasiliensis* kinase subfamily control network.

**Table S19b**. Edge betweenness values calculated across the *H. brasiliensis* kinase subfamily stress network.

## Supplementary Figures

**Fig. S1**. Phylogenetic analysis of 2,842 *H. brasiliensis* putative typical kinase proteins with 1,000 bootstrap replicates. Each protein is represented by a unique branch tip, and branch colors represent the kinase subfamily classification.

**Fig. S2**. Phylogenetic analysis of 2,842 *H. brasiliensis*, 1,531 *M. esculenta*, and 863 *R. communis* putative typical kinase proteins with 1,000 bootstrap replicates. Each protein is represented by a unique branch tip, and branch colors represent the kinase subfamily classification.

**Fig. S3**. Chromosomal positions of (A) 2,842 PK genes identified in *H. brasiliensis* and 1,531 PK genes identified in *M. esculenta*.

**Fig. S4**. (A) Molecular weight and (B) isoelectric point (pI) distribution of *H. brasiliensis* (Hbr), *M. esculenta* (Mes), and *R. communis* (Rco) protein kinases.

**Fig. S5**. Subcellular localization predictions of (1) *H. brasiliensis*, (2) *M. esculenta*, and (3)

*R. communis* protein kinases using LOCALIZER software (A), CELLO software (B), and Gene Ontology terms (C).

**Fig. S6**. Gene Ontology (GO) categories (biological process) of 339 tandemly duplicated *H. brasiliensis* protein kinases.

**Fig. S7**. Gene Ontology (GO) categories (biological process) of (A) 2,842 *H. brasiliensis*

protein kinases (PKs), (B) 1,531 *M. esculenta* PKs, and (C) 863 *R. communis* PKs.

**Fig. S8**. RNA expression profiles of *H. brasiliensis* protein kinases among 42 samples under control conditions. Sample IDs (columns) and subfamily names (rows) were clustered based on Euclidean distances.

**Fig. S9**. RNA expression profiles of *H. brasiliensis* protein kinases among 37 samples under abiotic stress conditions (cold, drought, jasmonate and ethylene treatments). Sample IDs (bottom) and subfamily names (right) were clustered based on Euclidean distances.

**Fig. S10**. RNA expression profiles of *H. brasiliensis* PKs among all 79 samples under control and stress conditions.

## Notes

### Competing Interest Statement

The authors have declared no competing interest.

### Summary of Updates

Text changes.

