## Supplementary figures and images for "The rubber tree kinome: genome-wide characterization and insights into coexpression patterns associated with abiotic stress responses"

### FigS3.jpg

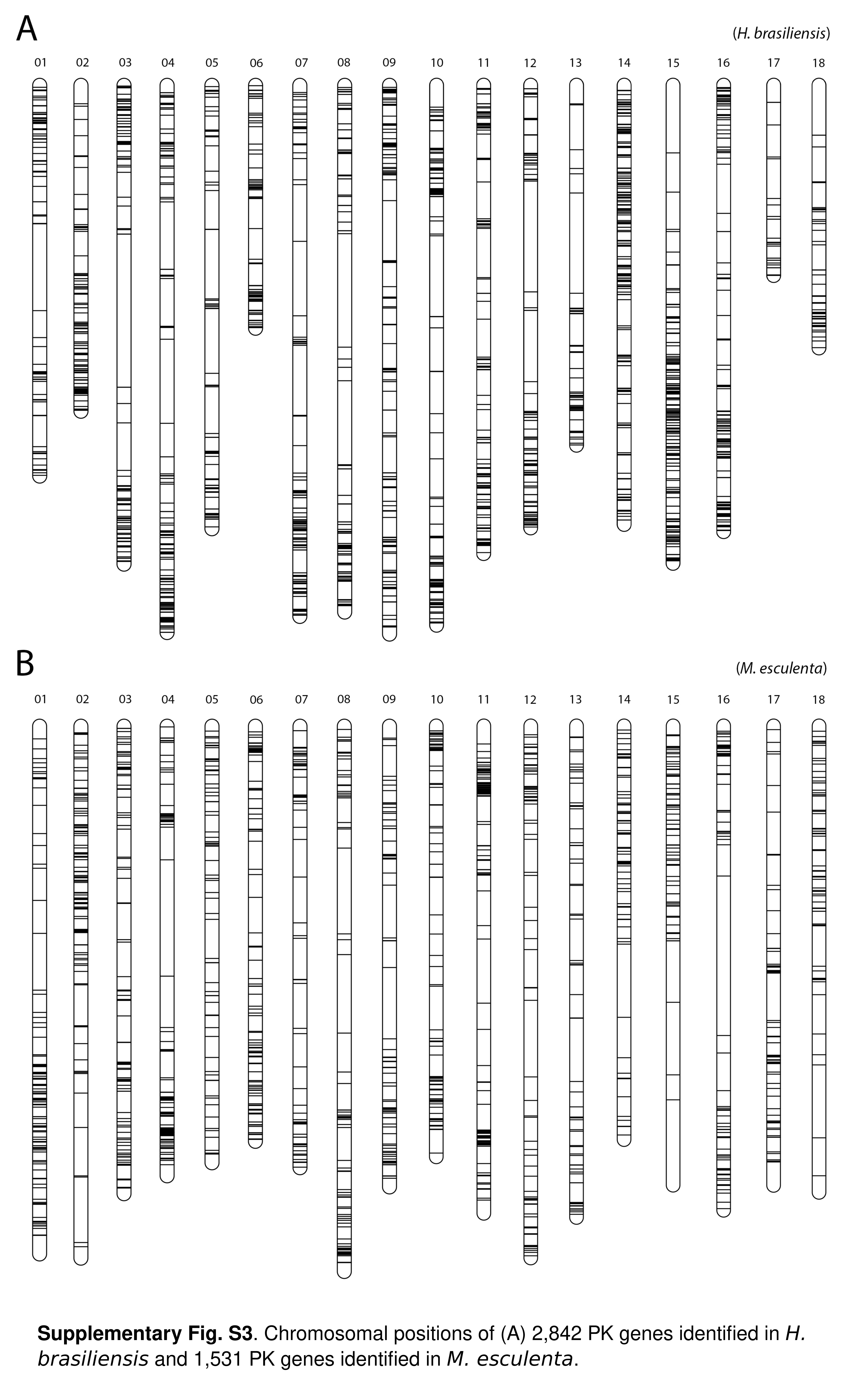

### FigS4.jpg

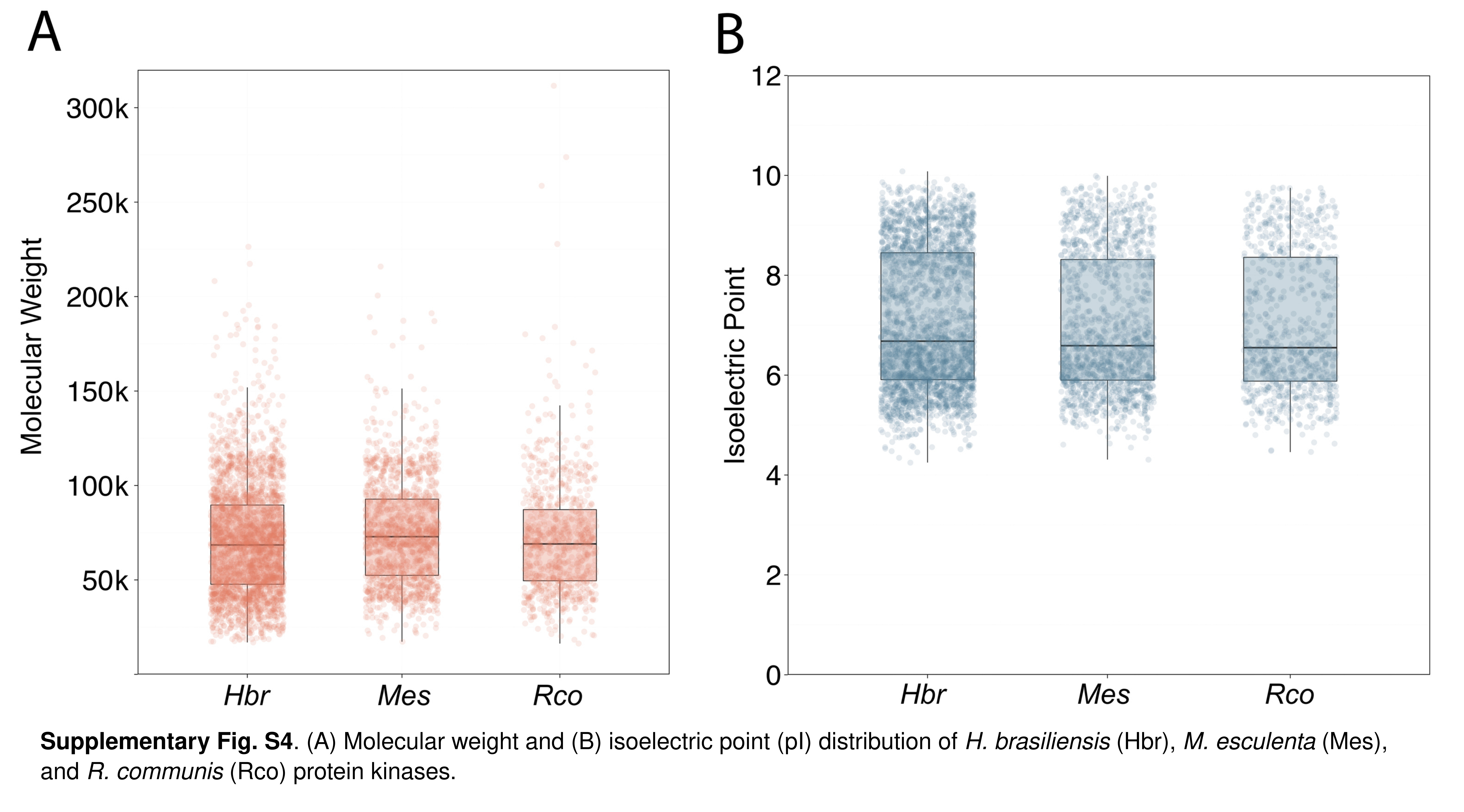

### FigS5.jpg

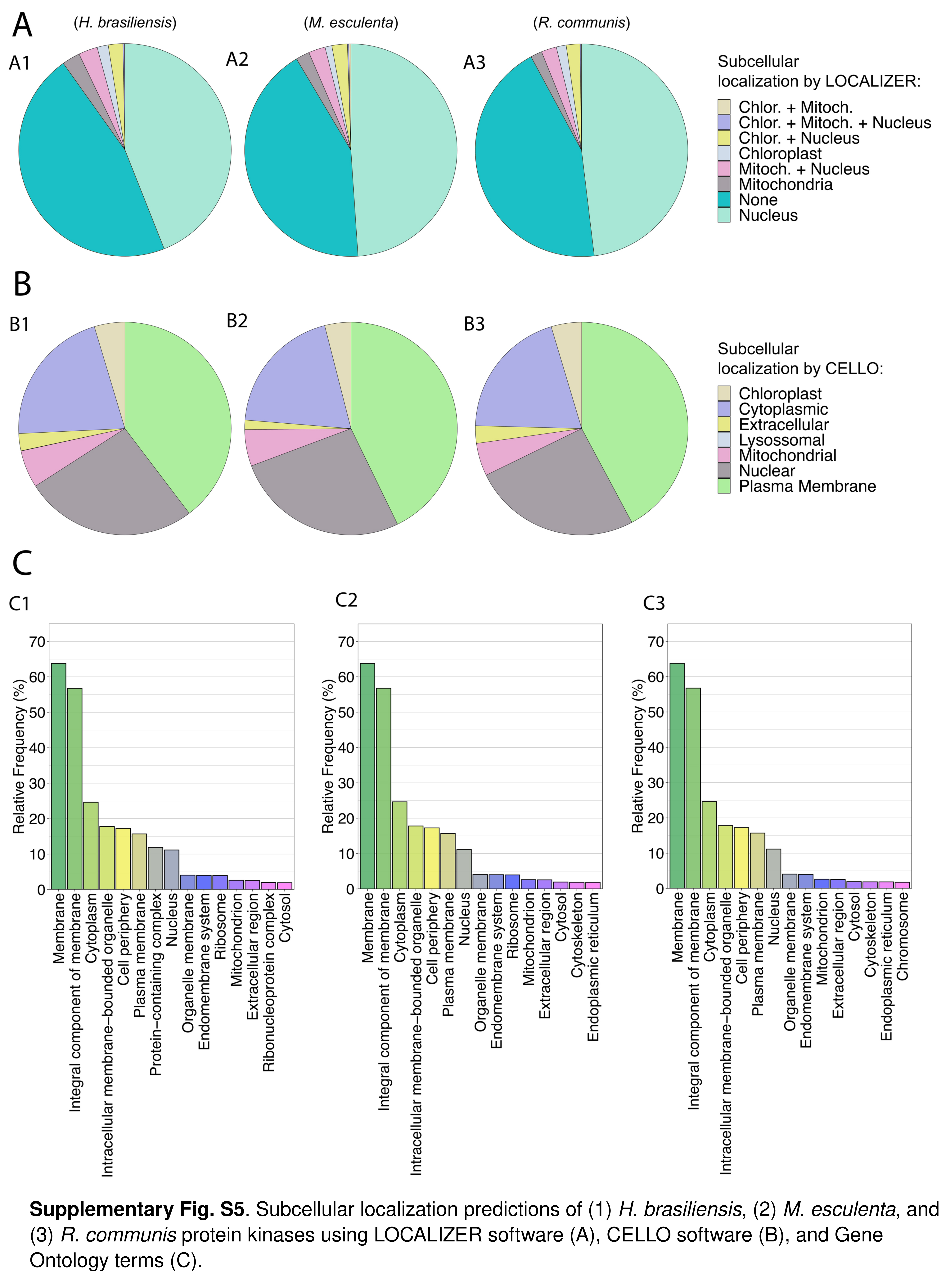

### FigS6.jpg

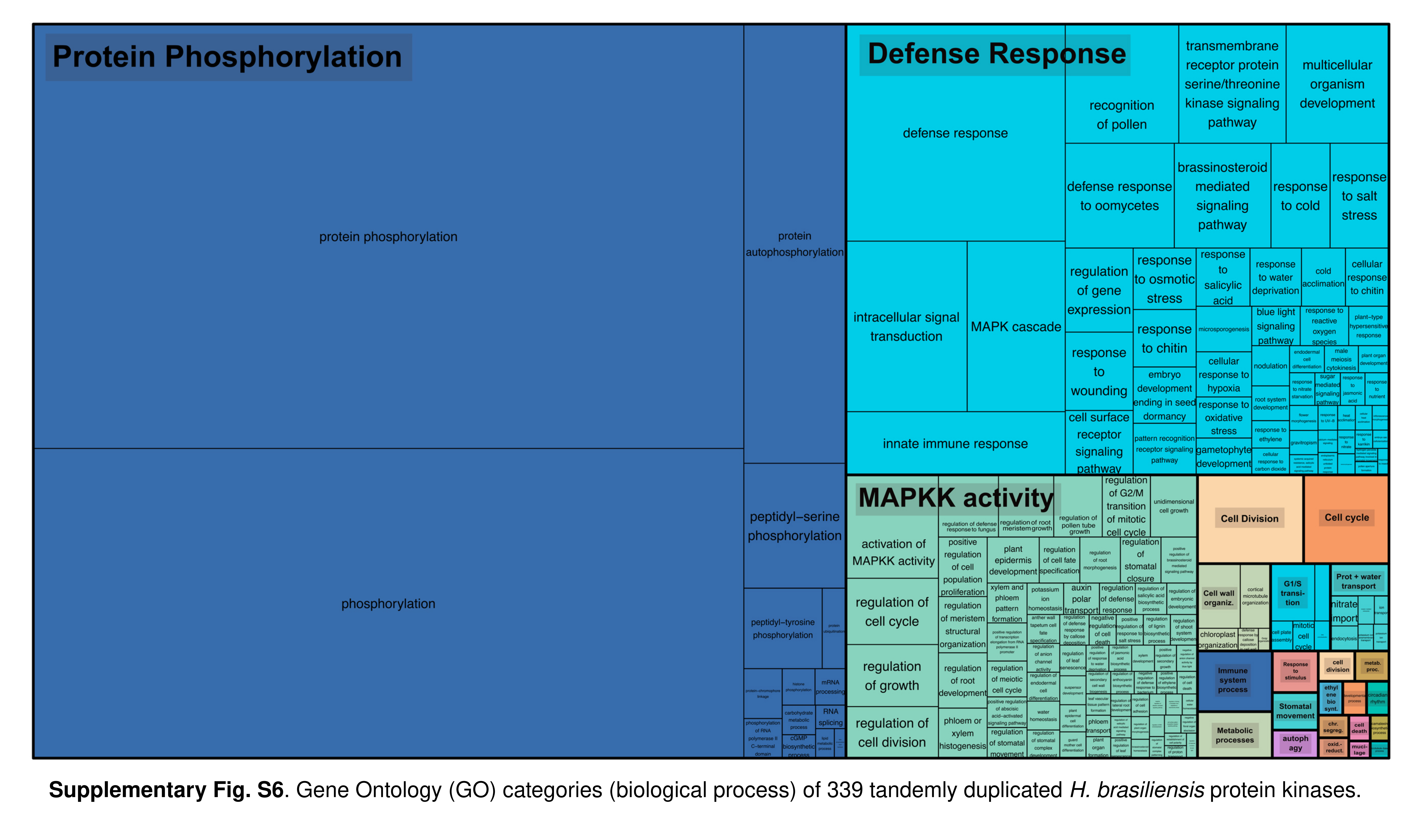

### FigS7.jpg

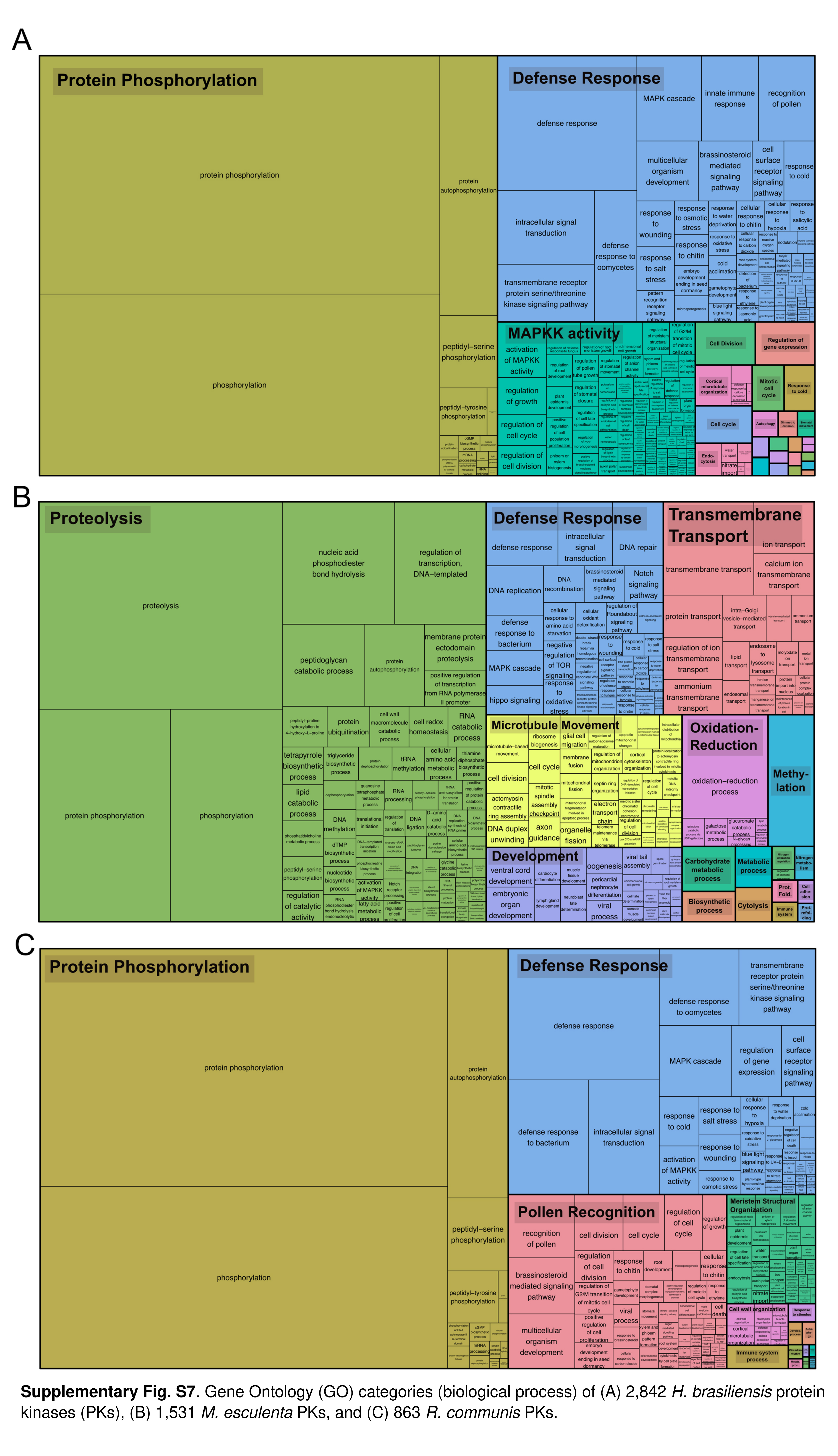

### FigS8.jpg

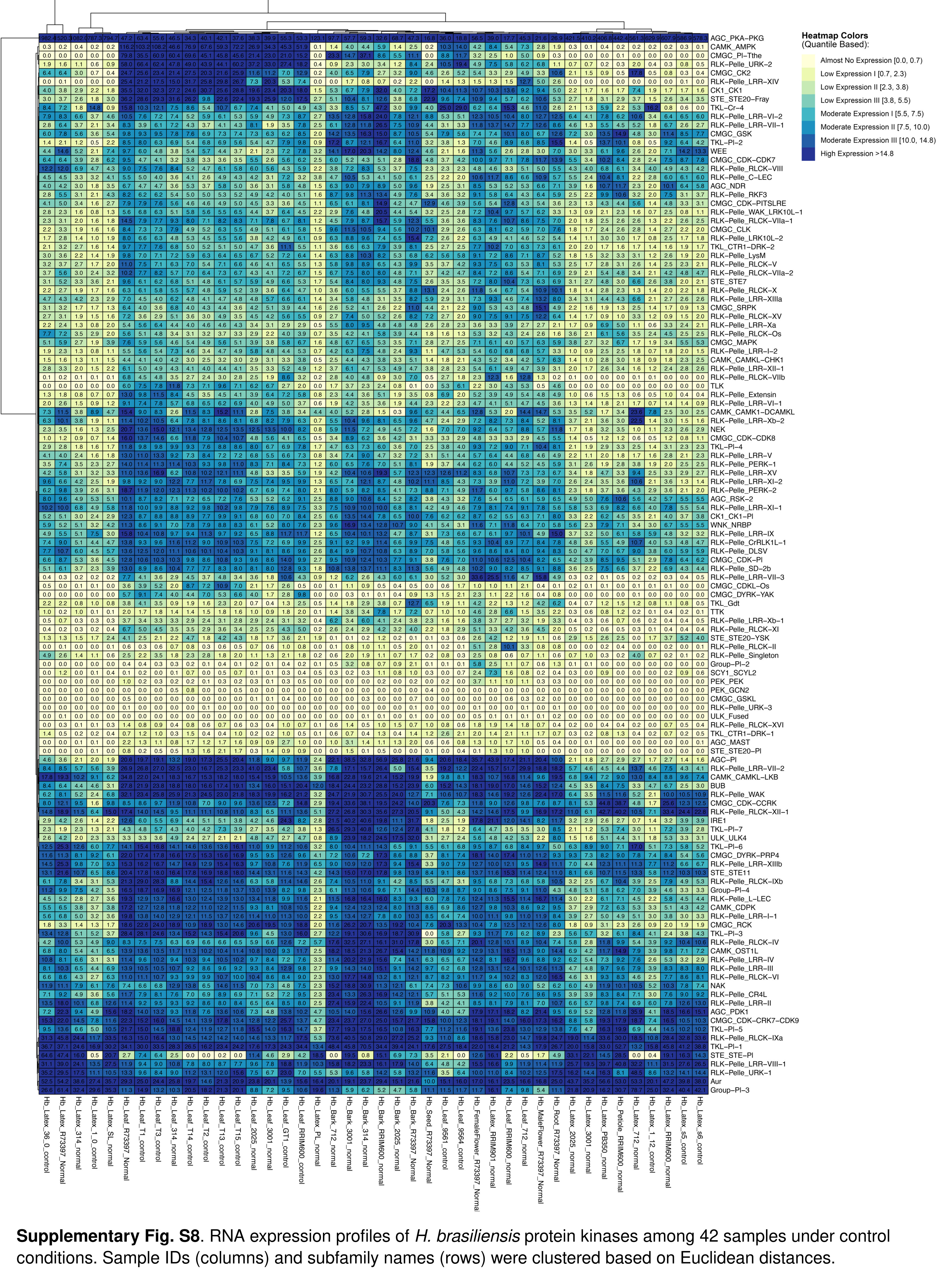

### FigS9.jpg

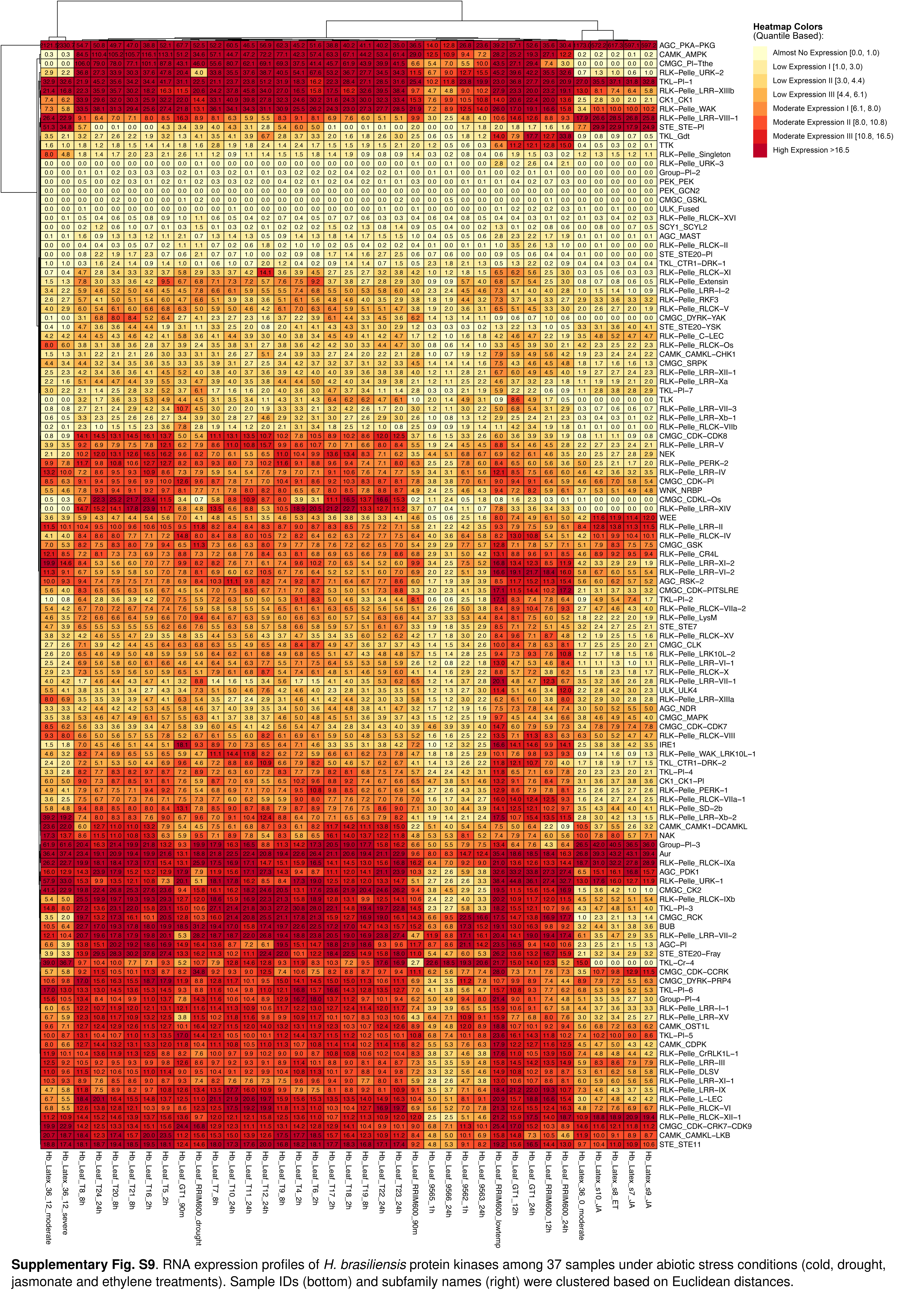

### FigS10.jpg

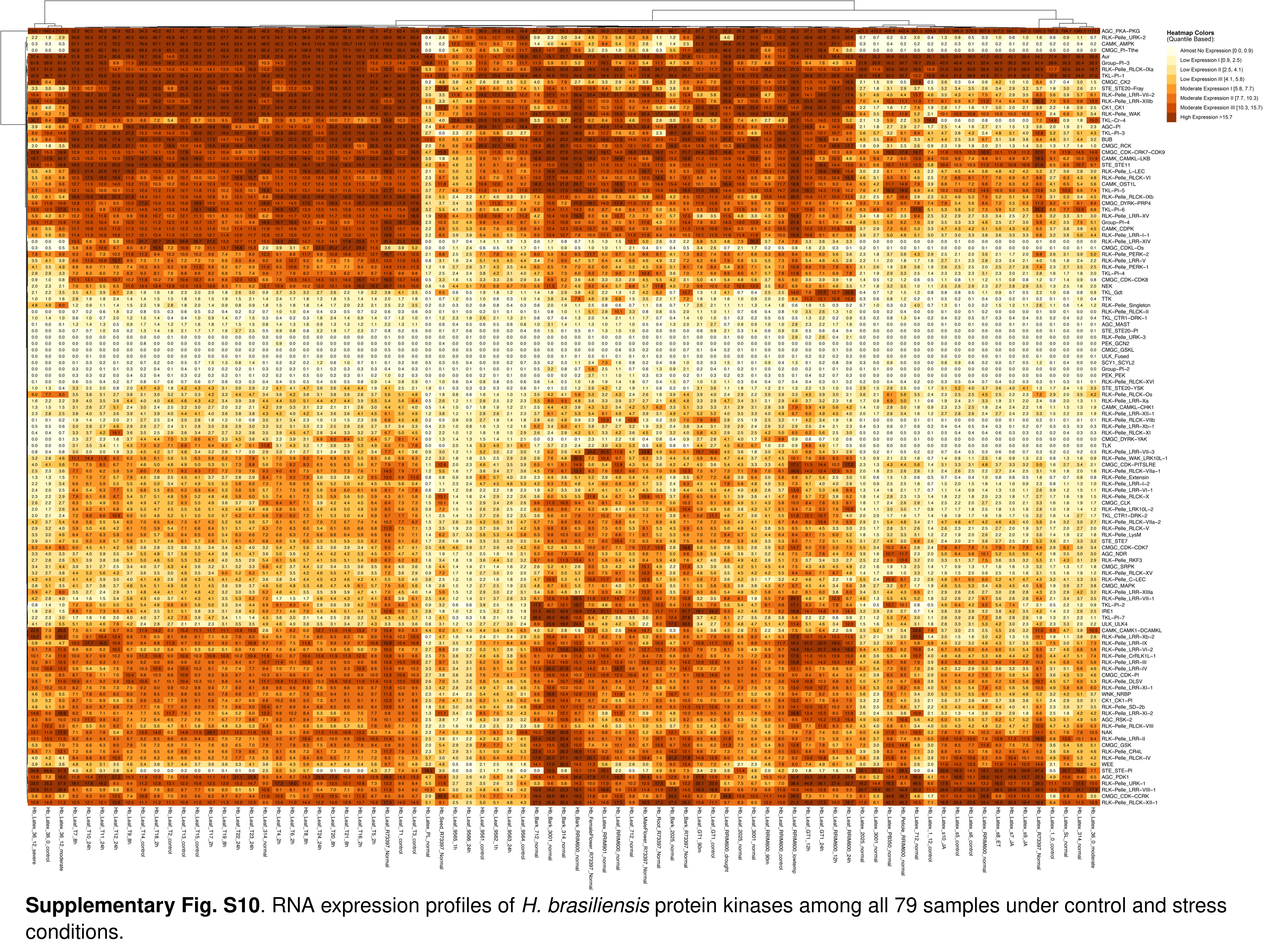
